# Decreased K_ATP_ channel activity contributes to the low glucose threshold for insulin secretion of rat neonatal islets

**DOI:** 10.1101/2021.03.04.433947

**Authors:** Juxiang Yang, Batoul Hammoud, Changhong Li, Abigail Ridler, Daphne Yau, Junil Kim, Kyoung-Jae Won, Charles A. Stanley, Toshinori Hoshi, Diana E. Stanescu

**Author notes:** the authors contributed equally to this work (Toshinori Hoshi, Diana E. Stanescu).

## Abstract

Transitional hypoglycemia in normal newborns occurs in the first 3 days of life and has clinical features consistent with hyperinsulinism. We found a lower threshold for glucose-stimulated insulin secretion from freshly isolated embryonic day (E)22 rat islets, which persisted into the first postnatal days. The threshold reached the adult level by postnatal day (P)14. Culturing P14 islets also decreased the glucose threshold. Freshly isolated P1 rat islets had a lower threshold for insulin secretion in response to BCH (2-aminobicyclo-(2,2,1)-heptane-2-carboxylic acid), a non-metabolizable leucine analog, and diminished insulin release in response to tolbutamide, an inhibitor of β-cell K_ATP_ channels. These findings suggested that decreased K_ATP_ channel function could be responsible for the lower glucose threshold for insulin secretion. Single-cell transcriptomic analysis did not reveal a lower expression of K_ATP_ subunit genes in E22 compared to P14 β-cells. The investigation of electrophysiological characteristics of dispersed β-cells showed that early neonatal and cultured cells had fewer functional K_ATP_ channels per unit membrane area. Our findings suggest that decreased surface density of K_ATP_ channels may contribute to the observed differences in glucose threshold for insulin release.

## Introduction

Transitional neonatal hypoglycemia occurs in healthy newborns in the first days of life. Plasma glucose concentrations reach a nadir by 2 hrs after birth, to a mean of 55-65 mg/dl (3-3.6 mmol/L), and increase to above 70 mg/dl (3.8 mmol/L) after the first 2-3 days, regardless of delayed or early feeding of neonates(1–4). Clinical features suggesting hyperinsulinemic hypoglycemia during the first postnatal day include hypoketonemia, non-suppressed plasma insulin concentrations, and inappropriately large glycemic response to glucagon or epinephrine (as reviewed in (5)). A similar timing of hypoglycemia occurs in rodent models, where insulin also appears to be secreted at low glucose concentrations postnatally (6, 7). While transitional neonatal hypoglycemia is typically mild, its underlying mechanism is unknown. Furthermore, there are few clinical criteria to differentiate this mild hypoglycemia from severe or persistent forms of hypoglycemia, which can cause irreversible brain injury and long-term neurologic defects.

Fetal plasma glucose concentrations closely follow maternal plasma concentrations(8). However, since maternal insulin does not cross the placenta, fetal β-cells are solely responsible for the fetal serum level of insulin, the major growth factor during the intrauterine period. If insulin secretion or signaling is impaired, fetal growth is restricted.(9, 10) If fetal β-cells have the same glucose threshold for glucose-stimulated insulin secretion (GSIS) as adult β-cells, fetal insulin secretion would be easily suppressed and fetal growth would be impaired. Hence, a β-cell adaptation that lowers the threshold for responding to glucose may be required to allow the fetus to secrete sufficient insulin to maintain growth. The clinical features of hyperinsulinemic transitional neonatal hypoglycemia are consistent with the concept that the low fetal glucose threshold for insulin secretion persists into the early neonatal period.

We aimed to characterize the glucose threshold for insulin secretion during the fetal and early neonatal period and to uncover the mechanism underlying the hyperinsulinism causing transitional neonatal hypoglycemia. We describe here the change in glucose threshold from late embryonic period to adulthood in rats and show that decreased K_ATP_ channel function, potentially through changes in K_ATP_ channel trafficking to the plasma membrane, is a major factor that contributes to the lower glucose threshold of embryonic and neonatal β-cells.

## Methods

### Animals

All animal experiments were performed in rats at the Childrens Hospital of Philadelphia with IACUC approval. Pregnant Sprague Dawley rats at embryonic day (E)15 or E17 were purchased from Charles River Laboratories (Wilmington, MA). Dams and pups were housed together in AAALC approved rodent colony. Male and female pups were batched together for measurements. Plasma glucose concentrations were determined using a Contour Next glucose meter (Ascensia Diabetes Care, NJ).

### Islet isolation and islet perifusions

Islet isolation was performed as previously described (11). In brief, pancreases were collected at E22 (last day of gestation in rats), postnatal day (P)1, P3, P7, P14, and P28 rats of both sexes. To minimize the number of animals used for these studies, we chose to perform only limited studies in E22 embryos. Pancreases were digested with collagenase (Millipore Sigma, Burlington, MA) in Hanks buffer. Islets were handpicked twice and then incubated in regular islet media (RPMI 1640, supplemented with 5 mM glucose, 10% fetal bovine serum, 2 mM glutamine, 1% antibiotics (Gibco, Gaithersburg, MD)) at 37°C for 3 hrs or, in some experiments, for 1 or 2 days, as specified. Islets, 200-250 in number, isolated from 1-5 rat pups, depending on postnatal age, were perifused at a flow rate of 1 ml/min. Perifusion media was a Krebs-Ringer bicarbonate buffer supplemented with 0.25% bovine serum albumin and 2 mM glutamine. Because of the low rates of insulin secretion in fetal islets, perifusion media was also supplemented with 0.1 mM 3-isobutyl-1-methylxanthine (IBMX), a phosphodiesterase inhibitor, to augment Ca^2+^-stimulated insulin secretion and bring insulin concentrations to the detection range of the insulin assay. After 30 min of perifusion with glucose-free buffer, step-wise increasing concentrations of glucose, BCH (2-aminobicyclo-(2,2,1)-heptane-2-carboxylic acid) or tolbutamide were added, every 20 min, as described in the figures. For the perifusions with BCH or tolbutamide, the perifusion media contained no glucose for the entire duration of the experiment. These experiments with BCH or tolbutamide were performed only at P1 and P14 because these were the ages with the largest difference in glucose threshold. After return to basal conditions for another 20 min, islets were exposed to 30 mM KCl to determine maximal insulin release at the end of the perifusion. Samples were collected every min for insulin assay using Insulin High Range Kits (Cat# 62IN1PEG, RRID:AB_2890910, Cisbio Assays, Bedford, MA) following the manufacturer’s instructions. For representation in Figures 1–3, % insulin release per min in response to glucose, BCH or tolbutamide was calculated as percentage of maximal KCl-stimulated insulin release for each replicate in each condition. In the presence of IBMX, maximal insulin secretion (ng/200-250 islets/min) in response to glucose and KCl, was approximately 30% lower in e22 islets compared to P1-P14 islets. Data points are connected with straight lines for presentation clarity only.

**Figure 1:**
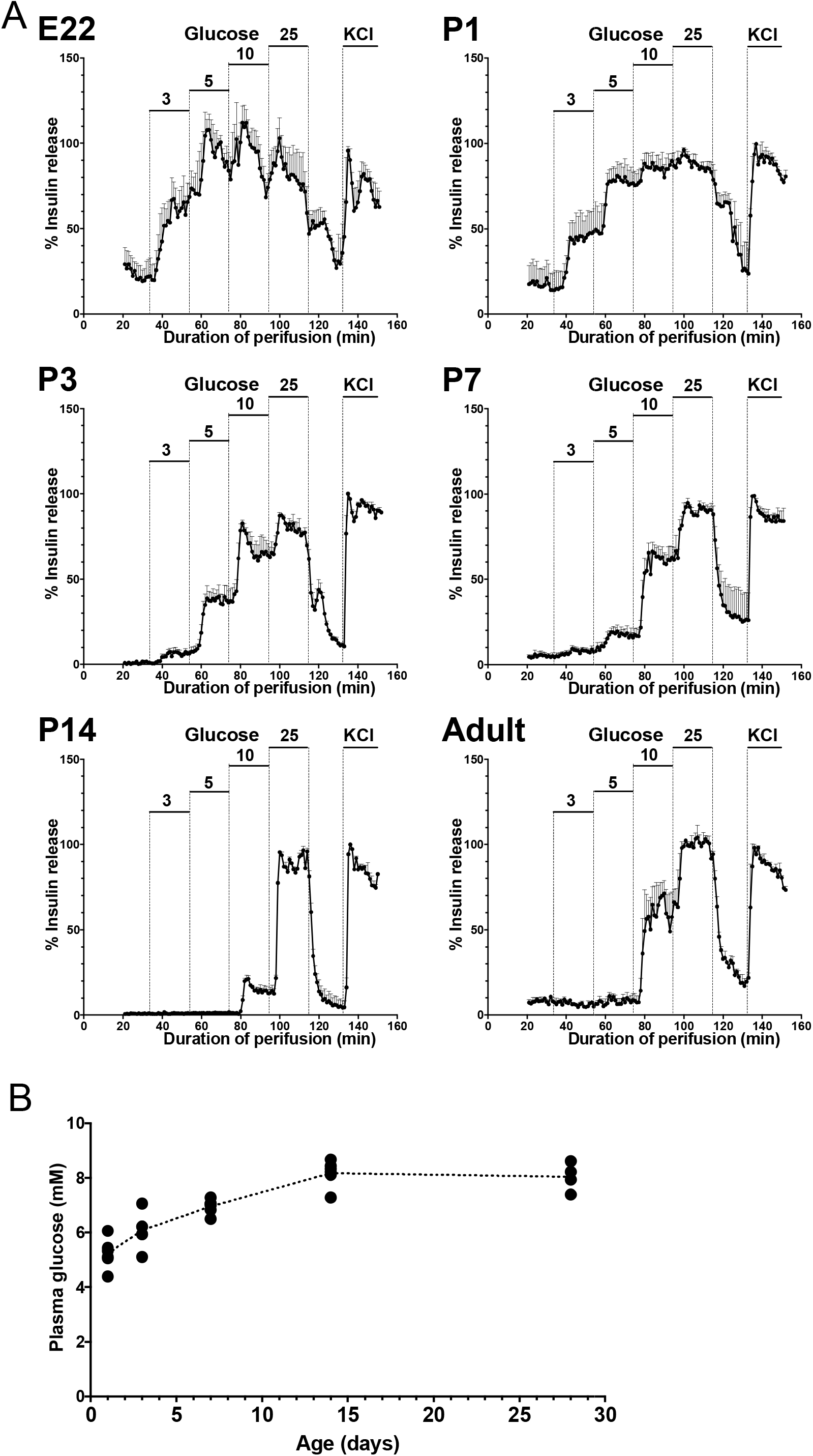
The glucose threshold for GSIS increases in early postnatal rat islets. **A.** Islet perifusions with stepwise increases in glucose concentration from 3 to 25 mM followed by 30 mM KCl for each age as depicted. Insulin release per min is calculated as percentage of maximal KCl-stimulated insulin release for each replicate. A total of 3-5 independent pools of freshly isolated islets (200-250 islets each) were obtained from 1-5 animals for each age. **B.** Plasma glucose concentrations of rat pups between 1-28 days of age. Each point represents measurements of one pup. Error bars in **A** and **B** represent SEM.

### Single-cell sequencing and analysis

Islets were isolated as described above from E22 embryos and P14 pups of both sexes. Islets were washed once with PBS without Mg^2+^ and Ca^2+^, followed by dissociation with TrypLe (Gibco, Gaithersburg, MD) for 7-9 min with gentle pipetting every 1 min. Dissociation was stopped by adding fetal bovine serum. The cell mixture was immediately filtered through 40 μm cell strainer, washed three times with cold PBS, and resuspended in regular islet media (as above). Single-cell suspensions were visualized under a microscope to assess cell viability and to ensure an adequate concentration for single-cell preparation. Cells were loaded on the 10X Genomics Chromium Controller (Pleasanton, CA) in the Center for Applied Genomics at CHOP, as per manufacturer’s instructions. Single-cell transcriptome libraries were sequenced on the HiSeq platform (Illumina, San Diego, CA). The CellRanger (v.2.1,10x genomics) pipeline was used for barcode filtering, alignment to *Rattus norvegicus* genome and Unique Molecular Identifiers (UMI) counting. Sequenced 10x libraries were individually evaluated using multiple criteria to determine cells of interest, including gene number (200 – 3000), number of UMI (0 – 20000), and percent mitochondrial gene expression (<0.10). The top 10 principal components were used to perform clustering and visualization using a t-distributed stochastic neighbor embedding (tSNE) plot. Gene marker expression was used to identify specific populations. We sequenced a total of 3556 cells, of which we identified 1072 cells (237 from E22 embryos and 835 from P14 pups) with transcriptomic signatures expected of β-cells (*Ins* and *Pdx1* expression). List of differentially expressed genes between E22 and P14 β-cells is provided in Supplemental Table (12).

### Whole-cell patch-clamp measurements

Electrophysiological measurements were performed using the apparatus previously described (13). Membrane potentials (V_m_) were monitored using the perforated whole-cell patch-clamp method with β-escin (MP Biomedicals, Santa Ana, CA) as the perforating agent (6 to 8 μM). The extracellular solution contained (in mM): 135 NaCl, 4 KCl, 2 CaCl_2_, 2 MgCl_2_, 10 mannitol, 5 glucose, 10 HEPES, pH 7.2 at 35°C with *N*-methyl-*D*-glucamine (NMG). The intracellular solution contained (in mM): 76 K_2_SO4, 10 KCl, 10 NaCl, 6 MgCl_2_, 30 mannitol, 30 HEPES, pH 7.2 at 35°C with NMG. Accounting for the divalent cation-chelating action of SO_4_^2-^, the free Mg^2+^ concentration is estimated to be ~2 mM. The tip of the wax-coated electrode (Warner G85150T, Holliston, MA) was filled with the intracellular solution and back-filled with the β-escin-containing intracellular solution. The current-clamp measurements were performed under a continuous perfusion condition (0.3 to 0.4 mL/min) at 35 °C (Warner TC-124A and TC-344B, Holliston, MA).

Voltage-clamp measurements were performed in the whole-cell configuration of the patch-clamp method at room temperature. Wax-coated patch electrodes (Warner G85150T) had a typical initial input resistance of 2 to 4 Mohms using the solutions described below. The external bath solution contained (in mM): 140 KCl, 2 MgCl_2_, 10 HEPES, pH 7.2 with *N*-methyl-*D*-glucamine (NMG). The internal pipette solution contained (in mM): 140 KCl, 11 EGTA, 10 HEPES, pH 7.2 with NMG. The absence of Mg^2+^ in the intracellular solution is not expected to impact activation of K_ATP_ channels without ATP (14). Once a gigaohm seal was obtained, the cell was transferred to a small perfusion chamber, ~150 μl in size, and then the membrane within the patch electrode was ruptured by negative pressure to obtain the whole-cell configuration. The cell was held at 0 mV and short 10 mV-square pulses were first applied to estimate total membrane capacitance and input resistance. Following electronic compensation for the capacitance and series resistance (~60%), voltage ramps from –80 mV to 80 mV in 400 ms were applied every 12 s. After establishing the whole-configuration, currents often increased in size, presumably because of loss intracellular components such as ATP, and the currents stabilized within 4 to 5 min, after which glyburide (400 nM; Sigma), a K_ATP_ channel inhibitor, was applied. Once the currents were stable in size in the presence of glyburide, the broad-spectrum K^+^ channel inhibitor quinidine (200 μM) was added to assess the gigaohm seal integrity. In some experiments, currents were recorded for up to 12 min before glyburide was applied to ensure that there was no slow change in current size.

The dissociated cells were amenable to electrophysiological measurements for up to 2 hrs following the cell isolation procedure. All salts were from Sigma (MilliporeSigma, Burlington, MA). The V_m_ values shown have been corrected for the junction potential.

## Results

### Glucose threshold for insulin secretion is lower in fetal and early postnatal islets

Figure 1A compares the insulin response of islets from E22 embryos, P1, P3, P7 and P14 pups, and adult rats to stepwise changes in glucose concentration to 3, 5, 10, and 25 mM. Glucose threshold was defined as the lowest glucose concentration examined at which insulin release exceeded 10% of the maximal release by 30 mM KCl above basal. The E22 islets showed a glucose threshold at 3 mM glucose. P1-P7 islets exhibited a progressive rightward shift in the glucose threshold with diminishing responses to 3 mM and then to 5 mM glucose; P14 and adult rat islets did not release insulin at 5 mM, but had a response to 10 mM glucose step, suggesting a glucose threshold between 5-10 mM. In E22 islets and at all other ages, the maximum insulin response to glucose in the presence of IBMX was similar to the response to membrane depolarization with KCl. The increase in threshold for GSIS closely paralleled the increase in plasma glucose concentrations between P1 to P14 (Fig. 1B). These observations suggest that that transitional neonatal hypoglycemia in human newborns is due to a lower fetal glucose threshold for GSIS, which is maintained in the immediate postnatal period (5).

### The low glucose threshold in neonatal islets reflects differences in the insulin secretory pathway distal to glycolysis

To assess whether the low glucose threshold for insulin release in fetal and newborn islets is conferred within the glycolytic pathway, we examined the responses of P1 and P14 islets to increasing concentrations of BCH, a non-metabolizable leucine analog. BCH promotes insulin release by activating β-cell mitochondrial glutamate dehydrogenase to stimulate oxidation of glutamate via α-ketoglutarate largely bypassing the glycolytic pathway, leading to a rise in β-cell ATP/ADP ratio and triggering insulin secretion (15). As shown in Fig. 2A, the BCH threshold for insulin secretion was lower in P1 islets (1 mM) compared to P14 islets (3 mM), suggesting that the low glucose threshold for insulin release in fetal and neonatal islets is determined, at least in part, at a step downstream of the glycolytic pathway.

**Figure 2:**
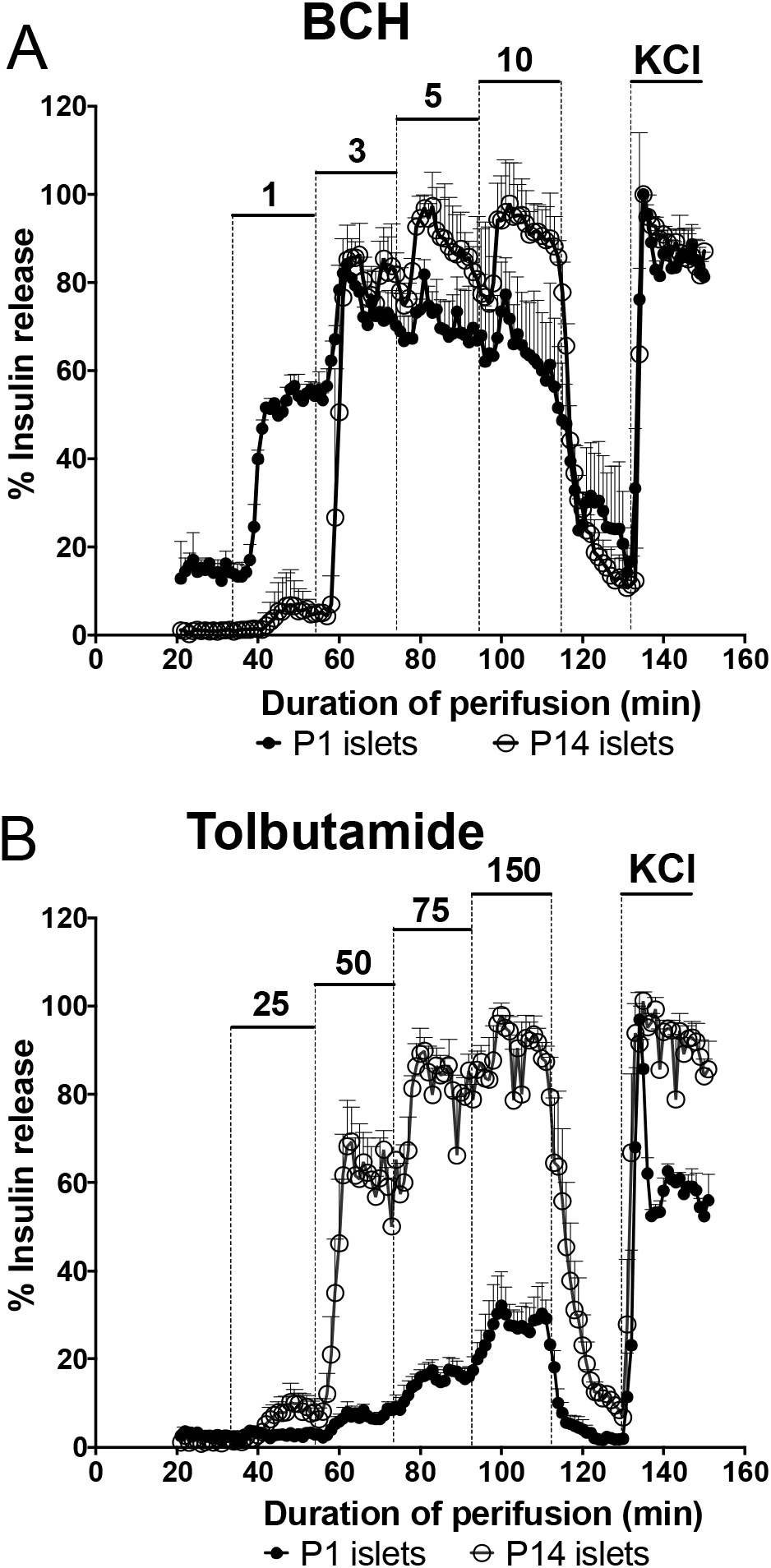
The lower threshold for insulin secretion is conferred by changes in the distal steps of insulin secretion pathway. **A:** Perifusions with stepwise increases in BCH concentration from 1 to 10 mM followed by 30 mM KCl in P1 and P14 freshly isolated rat islets. **B:** Perifusions with stepwise increases in tolbutamide concentration from 25 to 150 μM in freshly isolated P1 and P14 islets. Insulin release per min is calculated as percentages of maximal KCl-stimulated insulin secretion for each replicate and age. Three independent pools of islets (200-250 islets each) were obtained from 1-5 animals for each age. Error bars represent SEM.

To focus on the steps distal to mitochondrial ATP production, we assessed insulin responses to tolbutamide, an inhibitor of β-cell K_ATP_ channels that causes plasma membrane depolarization to trigger influx of Ca^2+^ and subsequent insulin release. The insulin response was markedly lower at all concentrations of tolbutamide in P1 islets compared to P14 islets, despite similar maximal insulin responses to KCl-induced depolarization (Fig 2B). This observation suggested that K_ATP_ channel activity and/or channel surface expression is lower in P1 than in P14 islets because a similar pattern of insulin secretion is seen in sulfonylurea receptor 1 (SUR1) knock-out islets, which have reduced insulin response to K_ATP_ channel inhibitors, but normal response to KCl (16–18).

### Culturing islets lowers the glucose threshold for insulin secretion

Davis and Matschinsky have previously shown that culturing human islets for 48 hrs decreased the glucose threshold for insulin secretion from 4-5 mM down to 2-3 mM (19). We confirmed this observation in rat P14 islets: culturing for 2 days decreased the glucose threshold from 10 mM to 3 mM (Fig. 3A). In contrast, for P1 islets, which already had a lower glucose threshold, 2 days of culture did not change the glucose threshold for insulin release (Fig. 3B). The insulin response to the K_ATP_ channel inhibitor tolbutamide was similarly diminished in P14 islets cultured for 2 days (Fig. 3C, D) and also in freshly isolated P1 islets (Fig. 2B, Fig. 3D). These findings show that P14 cultured islets have diminished responses to tolbutamide and implicate the changes in the K_ATP_ channels as the mechanism for the lower threshold for GSIS.

**Figure 3:**
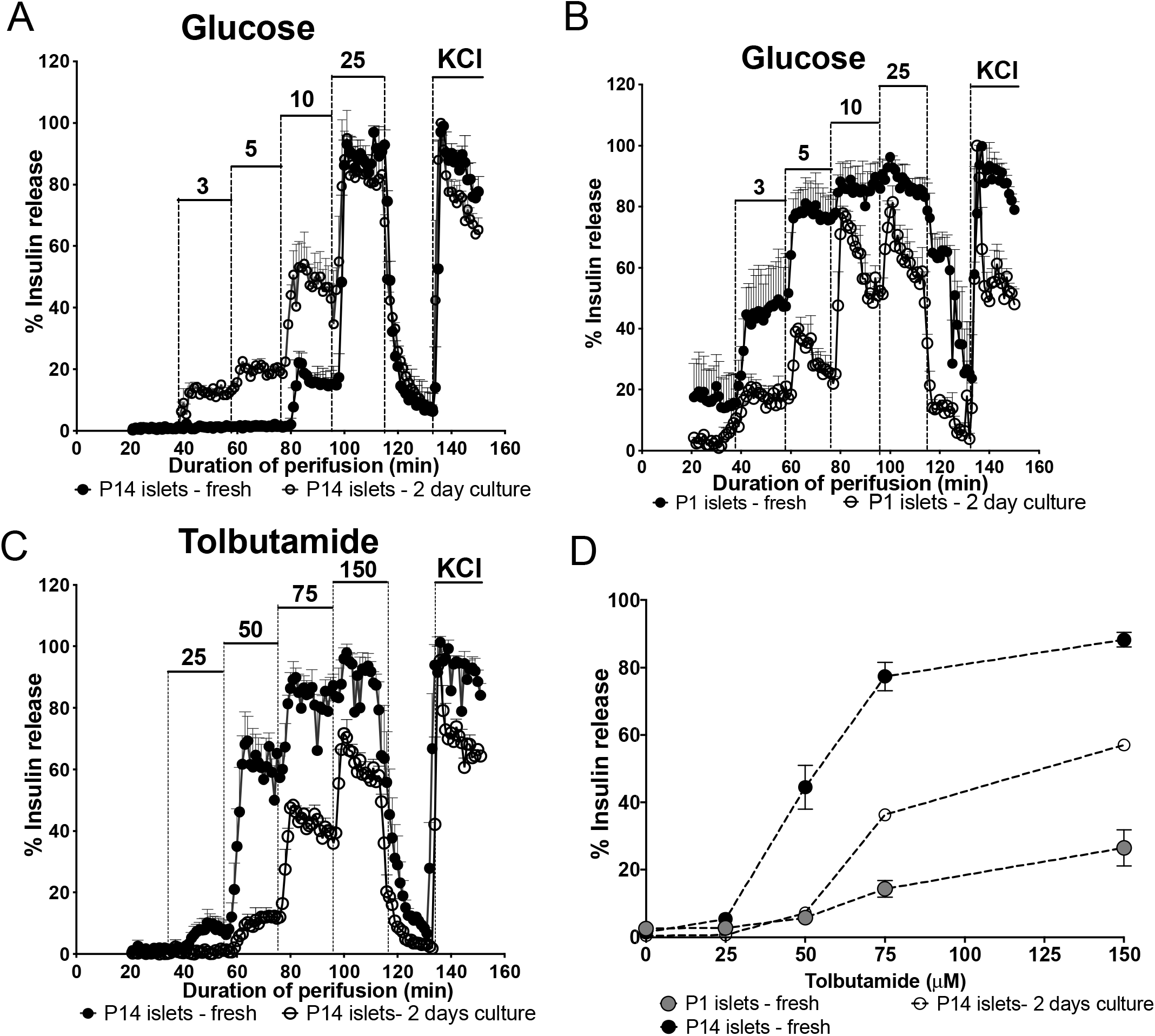
Glucose threshold for insulin release decreases after 2 days of culture. **A-B:** Perifusions with stepwise increases in glucose concentration from 3 to 25 mM followed by 30 mM KCl in freshly isolated P14 islets and P14 islets that had been cultured for 2 days (A) or freshly isolated P1 islets and P1 islets that had been cultured for 2 days (B). **C:** Perifusions with stepwise increases in tolbutamide concentration from 25 to 150 μM in freshly isolated islets and P14 islets that had been cultured for 2 days. **C:** Tolbutamide response curves in freshly isolated P1 islets, freshly isolated P14 islets, and cultured P14 islets. For **A-B**, 3-4 independent pools of islets (200-250 islets each) were obtained from 1-5 animals for each age. Insulin release per min is calculated as percentages of maximal KCl-stimulated insulin secretion for each replicate and age. Error bars represent SEM.

### Single-cell transcriptomic analysis of E22 and P14 cells

We performed differential expression analysis of single E22/P14 β-cells to identify gene transcript changes that could influence the threshold for GSIS. We identified 237 β-cells from E22 embryos and 835 β-cells from P14 pups (Fig. 4A-B); 135 transcripts were upregulated in E22 β-cells but only 17 transcripts were upregulated in P14 β-cells (false discovery rate <0.0001, logRatio>0.3) (Fig 4C; Suppl. Table) (12). β-cell maturational markers, such as *MafB, Npy* and *Nnat*, were upregulated in E22 β-cells compared to P14 β-cells, consistent with previous reports (6, 20, 21)(Fig. 4D). Among top upregulated genes in the P14 β-cells were *Sfrp5, Syt4, Ppy, Dbp*, and *Pcsk1n* (Fig. 4D, Suppl. Table) (12).

**Figure 4:**
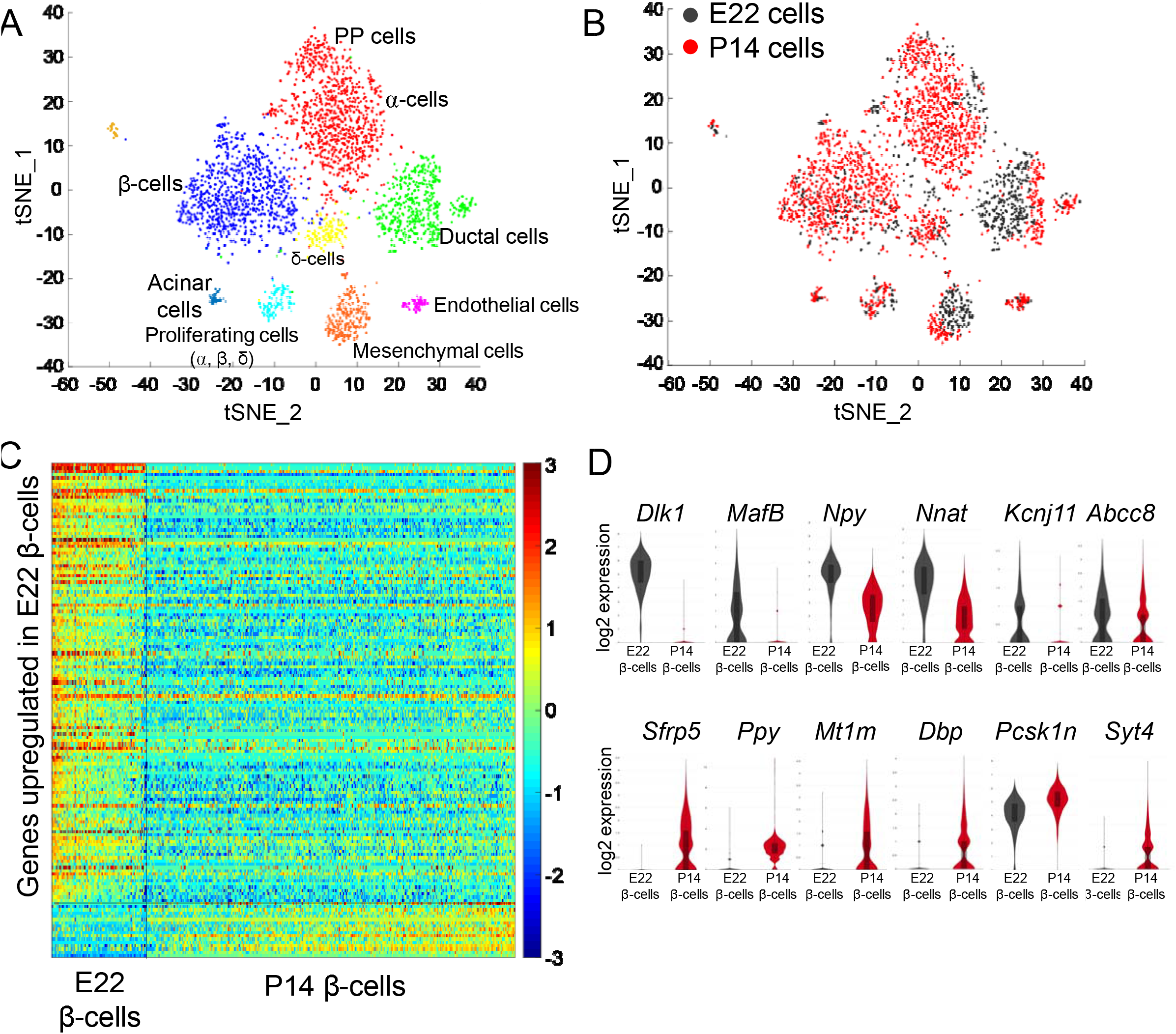
Single-cell transcriptomic analysis: **A-B.** t-SNE representation of E22/P14 isolated pancreas cells. **A:** Different clusters were identified based on expression of cell-specific genes. **B:** Cells are labeled depending on the age: E22 cells are presented in dark gray and P14 cells are represented in red. **C.** Heatmap of differentially expressed genes in E22 or P14 β-cells. **D:** Violin plots of log2 normalized expression levels (FPKM) for specific transcripts in E22 or P14 β-cells.

Several factors upstream of mitochondrial oxidation, including lactate dehydrogenase (*Ldh*), monocarboxylic transporter 1 (*Slc16a1*) and hexokinase isoforms 1-3 (*Hk1-3*), have been suggested to be responsible for the low glucose threshold for GSIS of early postnatal β-cells (22–24). These transcripts were not differentially expressed between the rat E22 and P14 β-cells. The lack of transcriptional changes, in the context of the perifusion data presented above, shows that the above factors are unlikely to be responsible for the low glucose threshold in fetal and neonatal islets.

The above results from the BCH-stimulated and tolbutamide-stimulated insulin secretion measurements suggested that decreased activity or expression of K_ATP_ channels, comprised of K_ir_6.2 channels complexed with SUR1, could be responsible for the lower glucose threshold of fetal and neonatal β-cells (see Fig. 2B). However, the mRNA level of *Abcc8* (SUR1) was higher in E22 β-cells and no difference in *Kcnj11* transcript (K_ir_6.2) was detected (Fig. 4D). These transcriptomic findings suggested that decreased transcription of K_ATP_ channel subunit genes is not the cause of the lower glucose threshold in neonatal islets.

### Neonatal islet cells have fewer functional K_ATP_ channels

The β-cell membrane potential (V_m_), in part controlled by K_ATP_ channels, is a critical regulator of insulin release (25). The lower glucose threshold in the fetal and early neonatal islets suggested the possibility that these β-cells have a V_m_ closer to the level at which voltage-dependent Ca^2+^ channels open to allow Ca^2+^ influx and subsequent insulin release. To test this prediction, we measured V_m_ values in individual cells from P3 and P14 islets incubated with 5 mM glucose. These two age groups were selected because the P3 islets were readily identified and amenable to enzymatic dissociation, whereas islets from younger animals were difficult to dissociate while preserving their functionality. In addition, as shown in Fig. 1A, at 5 mM glucose, P3 islets exhibited ~40% of maximal insulin release while P14 islets essentially showed no insulin release. V_m_ traces recorded from 29 cells isolated from 6 P3 pups and 33 cells from 9 P14 pups are illustrated in Fig. 5A. The V_m_ values show a high degree of variability in each group, ranging from near –90 mV to –30 mV (Fig. 5A; Suppl. Fig. 1) (12). Many of the cells exhibited non-overshooting action potentials (APs) typical of β-cells (Suppl. Fig. 1)(12). We observed that 41% of P3 cells had V_m_ values ≥–60 mV compared with 18% of P14 cells (Fig. 5A, B). This greater fraction of depolarized cells in the P3 group could contribute to the lower glucose threshold for GSIS found in the younger islets. While we do not have clear information regarding the cell-type identity of each cell in our electrophysiological measurements, in those experiments in which the glucose concentration was increased from 5 to 25 mM, the cells responded by firing non-overshooting APs, as expected from functionally viable β-cells (Suppl. Fig. 1) (12).

**Figure 5:**
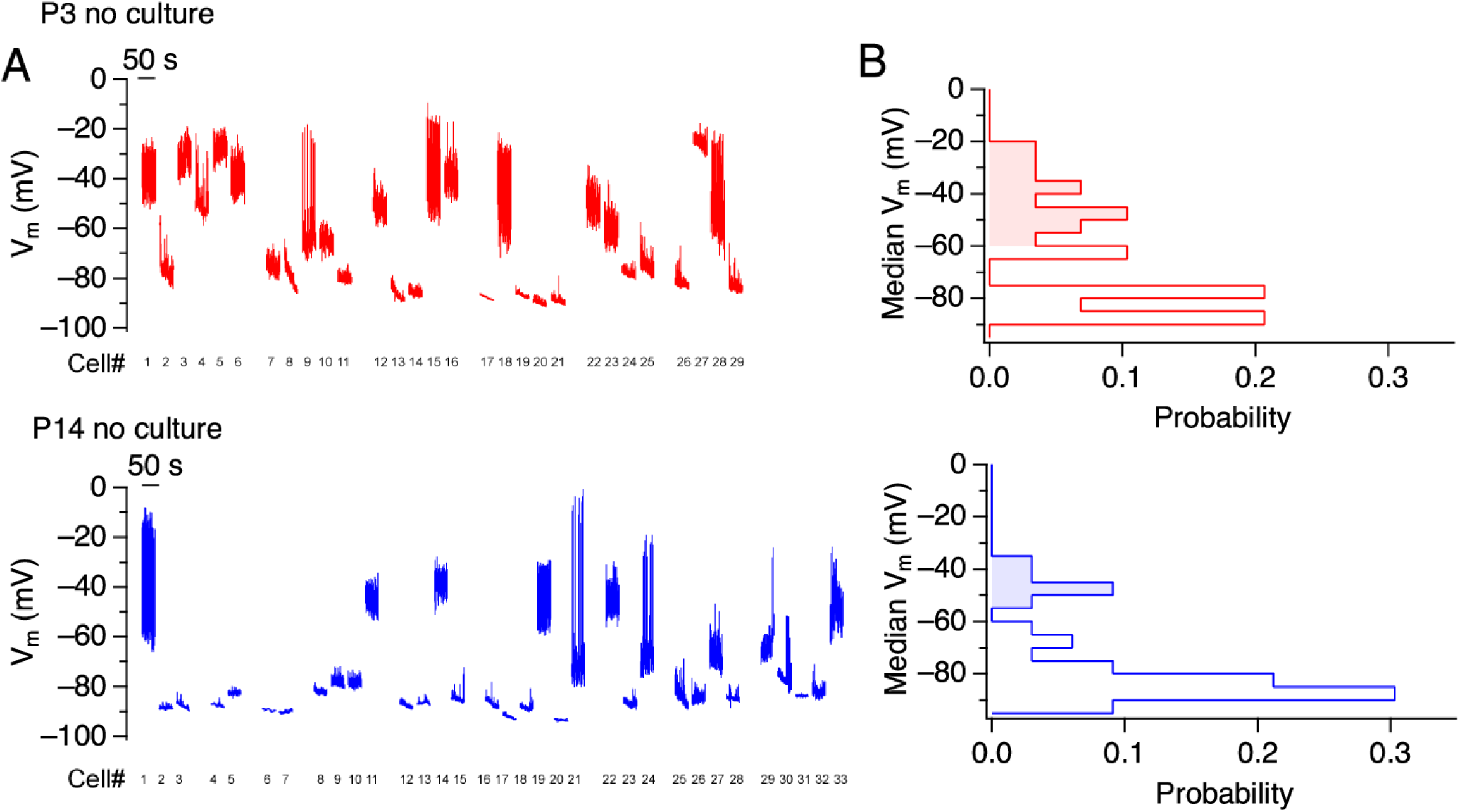
Membrane potential differences between P3 and P14 rat islet cells. **A-B** P3 rat cells have a more depolarized membrane potential (V_m_) at 5 mM glucose compared with P14 cells**. A**. Compressed 60-sec recordings of V_m_ values for from all cells measured: 29 P3 islet cells (red) and 33 P14 islet cells (blue). **B**. Probability density functions of median V_m_ values from the cells in P3 cells (red) and P14 cells (blue).

Plasma membrane depolarization in β-cells may be facilitated by closure/inhibition of K_ATP_ channels and/or a reduced number of channels trafficked to the cell surface (25, 26). Our perifusion results suggested that reduced K_ATP_ channel number or activity could contribute to the observed differences in the glucose threshold, both between E22-P14 islets and with culturing islets for 1 or 2 days. Total whole-cell membrane currents under a symmetrical 140 mM KCl condition were recorded from 20 P3 uncultured neonatal islet cells, 13 P14 uncultured islet cells, and 20 P14 islet cells cultured for 2 days. Input capacitance, reflecting the membrane area, was similar for uncultured P3 and P14 cells, and slightly greater the P14 cultured group (Suppl. Fig. 2A) (12). After establishing the whole-cell configuration, currents in response to repeated voltage ramps were recorded and illustrative results from a P14 cell are depicted in Suppl. Fig. 1B (12). The whole-cell configuration, establishing a small conduit through the membrane, disrupts the cellular enzymatic machinery. Therefore, the ramp currents immediately following establishing the whole-cell configuration reflect the near physiological ion channel status and those recorded later in time represent very compromised biochemical conditions when the ATP content is very low (Suppl. Fig. 2B; Fig. 6A-B) (12). Glyburide (400 nM) was next applied to inhibit K_ATP_ channels (Suppl. Fig. 2B; Fig. 6C) (12) and then quinidine, a broad-spectrum K^+^ channel inhibitor, was applied to assess the gigaohm seal integrity (Suppl. Fig. 2B) (12). The currents immediately after achieving the whole-cell configuration (~30 s) were noticeably greater in the uncultured P14 cells than in uncultured P3 or cultured P14 cells (Fig 6A). In most of the cells, the currents increased in size with time after achieving the whole-cell configuration, especially in P14 cells, probably due to disruption of ATP supply that activated K_ATP_ channels (Fig. 6B). The currents were subsequently nearly entirely blocked by the K_ATP_ channel inhibitor glyburide (Fig. 6C), suggesting that K_ATP_ channels dominate the whole-cell currents.

**Figure 6:**
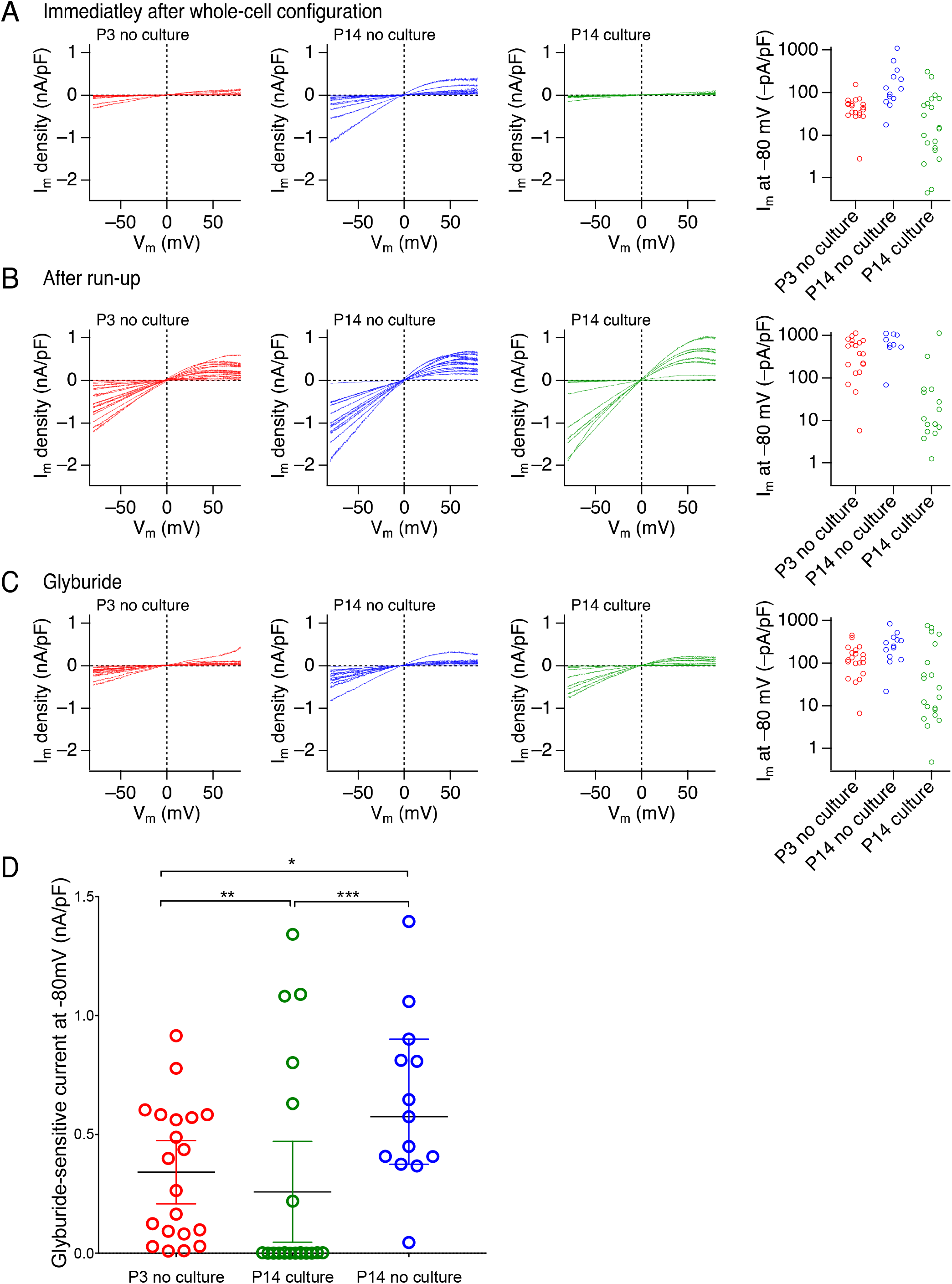
K^+^ currents in islet cells. Currents elicited by voltage ramps normalized by cell capacitance in individual cells **(A)** 20 to 30 s after whole-cell configuration, (**B**) after “run-up” of currents caused presumably by ATP loss and (**C**) after 400 nM glyburide. **D**. Freshly isolated P3 cells and cultured P14 cells have smaller glyburide-sensitive currents than uncultured P14 cells. Glyburide-sensitive currents normalized by cell capacitance at –80 mV for each cell measured. Each cell is represented by a circle. Results from uncultured P3 cells (*red*), uncultured P14 cells (*blue*) and cultured P14 cells (*green*). For A, B, and C, current sizes at –80 mV are shown (*right panels*). For D, statistical analysis: One Way ANOVA (Kruskal Wallis) p-value is 0.002; Mann-Whitney test for comparing two groups: (*) P3 no culture vs P14 no culture: p = 0.032, (**) P3 no culture vs P14 culture: p = 0.01, (***) P14 culture vs P14 no culture: p= 0.003. The p values are provided only as data description parameters.

Cultured P14 cells appeared to be comprised of two subpopulations based on the glyburide-sensitivity, with 60% (14 out of 20) of cultured P14 cells having no appreciable glyburide-sensitive currents (Fig. 6D – *green* circles). Uncultured P3 cells presented a similar two subpopulation distribution, with some of the cells exhibiting decreased glyburide-sensitive K^+^ currents compared to the uncultured P14 cells (Fig. 6D – *red* circles vs. *blue* circles). Thus, both P3 and cultured P14 islet cells had decreased glyburide-sensitive K^+^ currents, very likely K_ATP_ currents. These findings collectively suggest that decreased K_ATP_ channel activity, potentially due to decreased trafficking of channels to the plasma membrane, is the main contributor to the lower glucose threshold for GSIS in neonatal and cultured β-cells.

## Discussion

The results here demonstrate that fetal and early neonatal pancreatic β-cells respond to glucose stimulation with a lower glucose threshold than mature β-cells. This low threshold for GSIS correlates closely with the period of transitional hypoglycemia in normal newborns during the first day after birth. Our studies indicate that the glucose threshold for GSIS of early postnatal β-cells is lower than that of the adult β-cells at least until P7, and it reaches the adult value by P14. Furthermore, our results show that the differences in glucose threshold for GSIS during the early postnatal period reflect changes occurring distal to glycolysis and appear to involve the triggering pathway for insulin secretion at the site of the plasma membrane K_ATP_ channel. Electrophysiological measurements reveal reduced K_ATP_ channel activity, reflecting decreased density on the plasma membrane of neonatal β-cell. Culturing of isolated islets for 1-2 days produce a similar effect of reduced glucose threshold and reduced K_ATP_ activity, suggesting the possibility that extrinsic rather than developmental factors may be responsible for transitional neonatal hypoglycemia.

The repeated observation has been that isolated fetal β-cells are poorly responsive to glucose and only gain glucose responsiveness only after birth(27). Immature β-cells have been shown to have several differences in insulin secretory pathway when compared with mature β-cells: despite the presence of the insulin secretory machinery, it has been suggested there is a diminished coupling between glucose metabolism and membrane depolarization, leading to glucose insensitivity.(28–34) However, these observations from isolated islets must be carefully evaluated in the clinical context of β-cell function in the fetal and neonatal period to understand transitional hypoglycemia. Clinical evidence indicates that fetal β-cells are in fact responsive to glucose. Activating mutations in the glucose sensor of β-cells, glucokinase, lead to insulin-induced fetal overgrowth, whereas inactivating mutations in glucokinase lead to fetal undergrowth.(35–39) Increased fetal glucose levels due to maternal diabetes causes to greater fetal insulin secretion and large for gestational age birthweight (40). Hence, increased glucose phosphorylation in fetal β-cells increases insulin secretion, demonstrating that these cells have a functional insulin secretory pathway that is responsive to glucose, albeit with a different pattern of GSIS compared to adult β-cells.

A low glucose threshold for insulin secretion has been previously reported in both newborn rodent and human newborn islets (6, 41). The glucose threshold for GSIS, which determines the basal concentration of plasma glucose(19), is controlled by several components of the insulin secretory pathway, from the initial step of glucose phosphorylation to distal steps of insulin vesicle exocytosis. While glucokinase functions as the β-cell glucose sensor, mutations in many of the subsequent steps in the pathway of insulin secretion from glucokinase to the voltage-gated Ca^2+^ channel have been found to lead to congenital hyperinsulinism (42). The most common of the mutations alter K_ATP_ channel gene expression or activity, reflecting the key role of this channel in insulin regulation. Loss of function mutations in the K_ATP_ channel subunits, *KCNJ11* and *ABCC8*, lead to congenital hyperinsulinemic hypoglycemia due to a failure to appropriately suppress insulin release (43, 44). Decreased gene expression of the K_ATP_ channel subunits has also been proposed to be responsible for the infantile hyperinsulinemic hypoglycemia and subsequent maturity onset diabetes associated with inactivating mutations of HNF1A and HNF4A transcription factors (MODY1 and MODY3) (45–48). Recently, the decreased insulin secretion induced by leptin has been shown to involve increased density of K_ATP_ channels on the plasma membrane surface caused by increased trafficking of channels (49). Leptin increases surface expression of the K_ATP_ channel through upregulation of AMPK and PKA pathways without affecting channel gating properties (50). A defect in this leptin response pathway in *Phpt1^-/-^* mice causes lethal neonatal hypoglycemia due to impaired trafficking and decreased surface density of K_ATP_ channels.(26)

Multiple mechanisms have been suggested to explain the lower glucose threshold of neonatal islets including increased expression of genes normally disallowed in mature β-cells (*Slc16a1, LDH* genes and hexokinase isoforms with low affinity for glucose) or decreased expression of *Syt4* leading to increased sensitivity of insulin vesicles to Ca^2+^ (7, 22–24). It has also been suggested that the lower glucose threshold is involved in β-cell replication since it appears to correlate with the higher proportion of proliferating β-cells in the early postnatal period (51). We conclude here that the above mechanisms are unlikely to play a major role in determining the lower glucose threshold for GSIS in rat neonatal islets for the following two reasons. Firstly, the lower threshold for BCH-stimulated insulin secretion suggests that the low glucose threshold in fetal and neonatal islets is conferred at a step downstream of mitochondrial oxidation. This finding is in accord with the observed lack of transcript level changes in *Hk1-3, Slc16a1* or *Ldh* between E22 and P14 β-cells. Secondly, we have shown that *Syt4* expression is lower in E22 compared to P14 β-cells, supporting the role of this factor in the postnatal β-cell functional changes(7). However, the resulting increase sensitivity of insulin granules to Ca^2+^ due to lower *Syt4* expression in fetal and early neonatal β-cells would predict an increased, not a decreased, tolbutamide-stimulated insulin secretion that we observed in early neonatal islets.

Our single-cell transcriptomic analysis failed to detect transcriptomic-level regulation of *Kcnj11* or *Abcc8* genes between E22 and P14, consistent with other similar single-cell or bulk transcriptomic data sets in mice (20, 33). This observation, together with the suppressed tolbutamide-induced insulin secretion, suggests that decreased K_ATP_ channel surface density in fetal and neonatal β-cells contributes to the low glucose threshold for GSIS, leading to transitional hypoglycemia in the early neonatal period. We thus postulate that the change we found in K_ATP_ current size is caused by decreased channel trafficking to plasma membrane.

Fetal and neonatal islets have poorer maximal insulin responses to glucose stimulation and a higher maximal response to amino acids compared to mature islets (6, 27, 28, 34). The experiments here did not attempt to address these contrasting maximal responses; however, our observation of reduced K_ATP_ channel activity may not only explain the low glucose threshold for GSIS of fetal islets but also their decreased maximal GSIS and increased responsiveness to amino acids. A similar decreased maximal GSIS and increased responsiveness to amino acids are seen in genetic deficiency of K_ATP_ channels in human infants and in rodents (11, 18, 52, 53). In islets with impaired K_ATP_ channels from both humans and rodents, the insulin-glucose response curve is flattened with incomplete suppression of insulin release at low glucose concentrations and diminished rise in insulin at high glucose concentrations(18). Human and mouse islets with K_ATP_ mutations also show an exaggerated response to a mixture of amino acids via the amplification pathway in the presence of elevated concentrations of cytosolic Ca^2+^ (18, 53).

A few limitations exist in our study reported here. First, we assessed dynamic insulin secretion from isolated islets, not in the context of the whole animal. The observed lower glucose threshold may reflect the possibility that β-cell function may be influenced by a different *milieu* of nutrient and other hormones. However, the change in glucose threshold documented here strikingly parallels the plasma glucose in the first 7-14 days of life (as shown in Fig. 1B). This tight correspondence suggests that the glucose threshold of islets *in vivo* is indeed similar to the measurements in the *ex vivo* setting. Second, while our results here show that the glucose threshold for GSIS from early postnatal β-cells is determined largely in the last steps of the insulin secretory pathway, we cannot rule out the contribution of changes in the proximal steps in this pathway. For example, differences in glucose phosphorylation, glycolysis and/or mitochondrial oxidation could contribute to the threshold and/or maximal stimulated insulin secretion. Third, our dynamic insulin measurements were done in the presence of IBMX, a phosphodiesterase inhibitor that increases cAMP levels and augments Ca^2+^-stimulated insulin secretion. Phosphodiesterase inhibitors augment both the first and second phase of insulin secretion from neonatal and infant islets but do not affect insulin secretion at subthreshold glucose concentrations (41, 54, 55). While dynamic insulin measurements from adult human islets do not typically require the addition of a phosphodiesterase inhibitor, such pharmacological agents are helpful for similar studies in early postnatal islets. Thus, in early postnatal β-cells the amplifying pathway of insulin secretion may not be fully developed, either because of differences in PKA pathway in β-cells and/or differences in inter-cellular connections with neighboring α-cells. Indeed, intercellular gap junctions or other cell-to-cell contacts can change the characteristics of insulin release (56, 57).

The lower glucose threshold for GSIS of cultured islets, previously observed in isolated islets that had been cultured for 48 hrs (19), can be explained by our finding here that 60% of cultured β-cells have markedly reduced glyburide-sensitive K^+^ currents. We note, however, that other factors such as changes in intracellular Ca^2+^ handling, could also contribute (58). The observation that fetal/neonatal islets and islets exposed to culture conditions share a similar low glucose threshold for GSIS suggests the possibility of similar underlying mechanisms driven by extrinsic factors. Both the interior of cultured islets and fetal islets have reduced oxygen and receive a relatively constant nutrient supply, as opposed to a pulsatile feeding pattern (59). More severe restrictions of fetal oxygen or nutrient supply, as in birth asphyxia or late intrauterine growth retardation, cause perinatal stress-induced hyperinsulinism that may persist for several weeks after delivery (60, 61). Mechanistically, changes in the mTOR/AMP-kinase (27) and/or the hypoxia-inducible factor pathway could signal the postnatal increase in K_ATP_ channel trafficking to the β-cell plasma membrane and increase the threshold for GSIS. Furthermore, the lower glucose threshold for GSIS in fetal and early postnatal β-cells may provide as of yet undiscovered adaptive advantages, particularly in view of the surge in counter-regulatory hormones such as cortisol in the early neonatal period.

In conclusion, we have established that fetal and early postnatal β-cell function are characterized by a low glucose threshold for GSIS. This low threshold correlates with decreased K_ATP_ channel surface density. These findings support the concept that transitional neonatal hypoglycemia represents an extension into the early postnatal period of the fetal pattern of insulin release from β-cells.

## Disclosure statement

all authors have nothing to disclose

## Acknowledgements and funding

We thank Dr. Franz Matschinsky and Dr. Show-Ling Shyng for critical review of the manuscript. We also thank the Center for Applied Genomics at The Children’s Hospital of Philadelphia. DES was supported by American Diabetes Association Junior Faculty Development Award (1-16-JDF086). CAS was supported by NIH grant R37-DK056268. TH was supported in part by DK098517.

## Data availability statement

All single cell transcriptomic datasets generated during and/or analyzed during the current study are not publicly available but are available from the corresponding author (DES) on reasonable request.

## Author contributions

JY, BH designed and performed experiments, interpreted the results with input from DES and wrote parts of the manuscript. CL designed and performed experiments, interpreted the results, contributed to the conceptual frame work. AR performed experiments and revised the manuscript. DY designed and performed experiments. JK performed the bioinformatic analysis, interpreted the data. KJW conceptualized the bioinformatic analysis and revised the manuscript. CAH conceptualized and supervised the project, wrote and edited the manuscript. TH conceptualized the electrophysiological approach, performed experiments, interpreted the data and wrote/edited the manuscript. DES conceptualized and supervised the project overall, analyzed the data, wrote/edited the manuscript.

## Notes

### Competing Interest Statement

The authors have declared no competing interest.

